# Neural event segmentation of continuous experience in human infants

**DOI:** 10.1101/2021.06.16.448755

**Authors:** Tristan S. Yates, Lena J. Skalaban, Cameron T. Ellis, Angelika J. Bracher, Christopher Baldassano, Nicholas B. Turk-Browne

## Abstract

How infants experience the world is fundamental to understanding their cognition and development. A key principle of adult experience is that, despite receiving continuous sensory input, we perceive this input as discrete events. Here we investigate such event segmentation in infants and how it differs from adults. Research on event cognition in infants often uses simplified tasks in which (adult) experimenters help solve the segmentation problem for infants by defining event boundaries or presenting discrete actions/vignettes. This presupposes which events are experienced by infants and leaves open questions about the principles governing infant segmentation. We take a different, data-driven approach by studying infant event segmentation of continuous input. We collected whole-brain fMRI data from awake infants (and adults, for comparison) watching a cartoon and used a hidden Markov model to identify event states in the brain. We quantified the existence, timescale, and organization of multiple event representations across brain regions. The adult brain exhibited a known hierarchical gradient of event timescales, from shorter events in early visual regions to longer events in later visual and associative regions. In contrast, the infant brain only represented longer events, even in early visual regions, with no timescale hierarchy. The boundaries defining these infant events only partially overlapped with boundaries defined from adult brain activity and behavioral judgments. These findings suggest that events are organized differently in infants, with longer timescales and more stable neural patterns, even in sensory regions. This may indicate greater temporal integration and reduced temporal precision during dynamic, naturalistic perception.

**Significance Statement:** Sensory input is continuous and yet humans perceive discrete events. This event segmentation has been studied in adults by asking them to indicate natural breaks in continuous input. This classic parsing task is impossible in infants who cannot understand or follow instructions. We circumvent this barrier by testing how the infant brain parses. We applied a computational model to rare awake fMRI data from infants to identify how their brains transitioned between stable states during a cartoon. Whereas adults showed a gradient in event timescales, from shorter events in sensory regions to longer events in associative regions, infants persistently segmented fewer, longer events across the cortical hierarchy. These findings provide new neuroscientific insight into how infants represent their environment.

## Introduction

From the moment we are born, our sensory systems are bombarded with information. Infants must make sense of this input in order to learn regularities in their environment ^[1,2]^ and remember objects and events ^[3,4]^. How infants overcome this perceptual challenge has consequences for other cognitive abilities, such as social competence and language ^[5]^. Adults segment continuous experience into meaningful events ^[6]^, both online ^[7,8]^ and after the fact^[9]^. This occurs automatically and at multiple timescales ^[10]^, allowing us to perceive the passage of long events (e.g., a talk from a visiting scientist) and to differentiate or integrate shorter events that comprise them (e.g., an impressive results slide or funny anecdote). These multiple timescales of event perception can be modulated by attentional states^[11]^ and goals^[10]^ to support adaptive decision-making and prediction^[12]^. Thus, characterizing event segmentation in infants and how it relates to that of adults is important for understanding infant perception and cognition.

Research on infant event perception has found that infants are sensitive to complex event types such as human actions^[2,13,14]^ and cartoons^[4,15]^. Infants recognize the similarity between target action segments and longer sequences that contain them, with greater sensitivity to discrete events (e.g., an object being occluded) than to transitions between events (e.g., an object sliding along the ground)^[16,17]^. Such findings suggest that infants not only segment experience at a basic sensory level, in reaction to changes in low-level properties, but are also capable of segmenting at a more abstract level like adults. Behavioral measures such as looking time have expanded our understanding of infant event processing. Yet, such measures provide indirect evidence and may reflect multiple different underlying cognitive processes. For example, the same amount of looking to a surprising event could reflect novelty detection or visual memory for components of the event ^[18]^. Thus, infant researchers are increasingly using neural measures such as electroencephalography (EEG) to study infant event processing. These studies find differences in brain activity to pauses that disrupt events versus coincide with event boundaries^[19–21]^. In sum, there is rich evidence across paradigms that infants are capable of event segmentation and that this guides their processing of ongoing experience.

However, a limitation in the current literature on infant event segmentation is that it relies on boundaries determined by (adult) experimenters. This reflects an unstated or unintended assumption that infants experience the same event boundaries as adults, which may obscure events that are specific to either age group. Because behavioral research with infants tests their looking preferences after part versus whole events ^[13,16]^, it would be infeasible to test all possible boundary locations and event durations. Brain imaging has the potential to contribute to our understanding by circumventing this limitation. Although most infant EEG studies have followed the behavioral work by introducing pauses at or between predetermined event boundaries, some adult EEG studies have found signatures of event segmentation during a continuous sequence of images ^[22]^ or a movie^[23]^. Although similar studies could be performed in infants, EEG may not have the spatial resolution or sensitivity away from the scalp to recover event representations in subcortical structures such as the hippocampus^[24]^ and midline regions such as the precuneus and medial prefrontal cortex^[25]^, each of which has been shown to play a distinct role in adult event segmentation^[26]^.

Functional magnetic resonance imaging (fMRI), has proven effective at capturing multiple event representations in adults during continuous, naturalistic experience ^[27]^. In one fMRI approach, behavioral boundaries from an explicit parsing task are used as event markers to model fMRI activity during passive movie watching. Regions such as the superior temporal sulcus and middle temporal area respond to events at different timescales, with larger responses at coarser boundaries^[8,28,29]^. Other brain regions respond transiently to different types of event changes, such as character changes in the precuneus and spatial changes in the parahippocampal gyrus^[8]^. However, applied to infants, this approach would suffer the aforementioned limitation of adult experimenters predetermining event boundaries. An alternative fMRI approach discovers events in a data-driven manner from brain activity^[25,30]^, using an unsupervised computational model to learn stable neural patterns during movie watching. This model can be fit to different brain regions to discover a range of event timescales. In adults, sensory regions process shorter events, whereas higher-order regions process longer events, mirroring the topography of temporal receptive windows ^[31–33]^. Moreover, event boundaries in the precuneus and posterior cingulate best match narrative changes in the movie^[25]^. Thus, fMRI could reveal fundamental aspects of infant event perception that cannot otherwise be accessed easily ^[34–38]^.

In this study, we collected movie-watching fMRI data from infants in their first year to investigate the early development of event perception during continuous, naturalistic experience. We also collected fMRI data from adults watching the same movie. We first asked whether the movie was processed reliably across infants using an intersubject correlation analysis^[39]^. Eye movements during movie-watching are less consistent in infants than adults^[40,41]^, and thus it was not a given that infant neural activity would be reliable during the movie. After establishing this reliability, we then asked whether and how infants segmented the movie into events using a data-driven computational model. As a comparison and to validate our methods, we first performed this analysis on adult data. We attempted to replicate previous work showing a hierarchical gradient of timescales in event processing across regions in the adult brain, with more/shorter events in early sensory regions at the bottom of the hierarchy and fewer/longer events in associative regions at the top of the hierarchy. This itself was an open question and extension of prior studies because the infant-friendly movie we used was animated rather than live action and was much shorter in length, reducing the range of possible event durations and the amount of data and statistical power.

With this adult comparison in hand, we tested three hypotheses about event segmentation in the infant brain. The first hypothesis is that infants possess an adult-like hierarchy of event timescales across the brain. This would fit with findings that aspects of adult brain function, including resting state networks^[42]^, are present early in infancy. The second hypothesis is that the infant brain shows a bias to segment events at shorter timescales than adults. This would result in a flatter hierarchy in which higher-order brain regions are structured into a greater number of events, akin to adult early visual cortex. Such a pattern would fit with findings that sensory regions mature earlier in development^[38]^, and may provide bottom-up input that dominates the function of (comparatively immature) higher-order regions. The third hypothesis is that the infant brain shows a bias to segment events at longer timescales than adults. This would also result in a flatter hierarchy, just in reverse, with fewer events in sensory regions, akin to higher-order regions in adults. This could reflect attentional limitations that reduce the number of sensory transients perceived by infants ^[43]^ or greater temporal integration in infant visual^[44,45]^ and multisensory processing^[46,47]^. We adjudicate these hypotheses and provide detailed comparisons between infant and adult event boundaries.

## Results

### Intersubject correlation reveals reliable neural responses in infants

We scanned infants (*N* = 24; 3.6–12.7 months) and adults (*N* = 24; 18–32 years) while they watched a short, silent movie (“Aeronaut”) that had a complete narrative arc. To investigate the consistency of infants’ neural responses during movie-watching, we performed leave-one-out intersubject correlation (ISC), in which the voxel activity of each individual participant was correlated with the average voxel activity of all other participants ^[39]^. This analysis was performed separately in adults and infants for every voxel in the brain and then averaged within eight regions of interest (ROIs), spanning from early visual cortex (EVC) to later visual regions (lateral occipital cortex, LOC), higher-order associative regions (angular gyrus, AG; posterior cingulate cortex, PCC; precuneus; medial prefrontal cortex, mPFC) and the hippocampus. Because the movie was silent, we used early auditory cortex (EAC) as a control region.

Whole-brain ISC was highest in visual regions in adults (Figure 1A), similar to prior studies with movies^[39,48]^. That said, all eight ROIs were statistically significant in adults (EVC: *M* = 0.498, CI = [0.446, 0.541], *p* < 0.001; LOC: *M* = 0.430, CI = [0.391, 0.466], *p* < 0.001; AG: *M* = 0.094, CI = [0.058, 0.127], *p* < 0.001; PCC: *M* = 0.143, CI = [0.097, 0.190], *p* < 0.001; precuneus: *M* = 0.160, CI = [0.128, 0.193], *p* < 0.001; mPFC: *M* = 0.053, CI = [0.032, 0.077], *p* < 0.001; hippocampus: *M* = 0.047, CI = [0.032, 0.062], *p* < 0.001; EAC: *M* = 0.081, CI = [0.044, 0.115], *p* < 0.001). Broad ISC in these regions is consistent with prior studies that used longer movies. The one surprise was EAC, given that the movie was silent (see Discussion for potential explanations).

**Figure 1.**
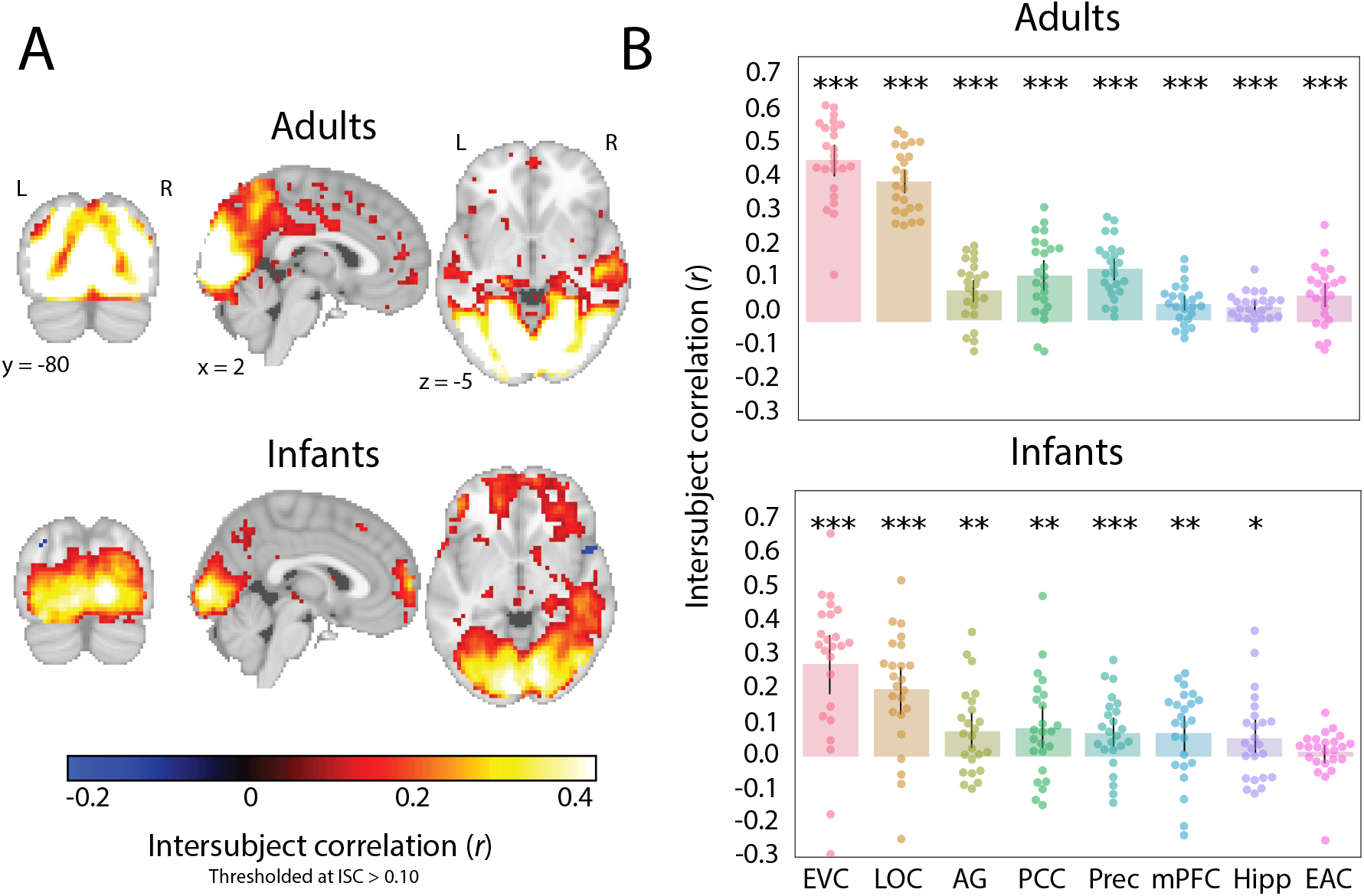
Average leave-one-out intersubject correlation (ISC) in adults and infants. (A) Voxel-wise ISC values in the two groups, thresholded arbitrarily at a mean correlation value of 0.10 to visualize the distribution across the whole brain. (B) ISC values were significant in both adults and infants across ROIs (except EAC in infants). Dots represent individual participants and error bars represent 95% CIs of the mean from bootstrap resampling. *** *p* < 0.001, ** *p* < 0.01, * *p* < 0.05, ∼*p* < 0.1. ROIs: early visual cortex (EVC), lateral occipital cortex (LOC), angular gyrus (AG), posterior cingulate cortex (PCC), precuneus (Prec), medial prefrontal cortex (mPFC), hippocampus (Hipp), early auditory cortex (EAC).

ISC was weaker overall in infants than adults, but again higher in visual regions compared to other regions. All ROIs except for EAC were statistically significant in infants (EVC: *M* = 0.290, CI = [0.197, 0.379], *p* < 0.001; LOC: *M* = 0.206, CI = [0.135, 0.275], *p* < 0.001; AG: *M* = 0.079, CI = [0.030, 0.133], *p* = 0.002; PCC: *M* = 0.087, CI = [0.028, 0.154], *p* = 0.006; precuneus: *M* = 0.073, CI = [0.030, 0.118], *p* < 0.001; mPFC: *M* = 0.073, CI = [0.017, 0.124], *p* = 0.009; hippocampus: *M* = 0.059, CI = [0.012, 0.113], *p* = 0.023; EAC: *M* = 0.013, CI = [-0.018, 0.039], *p* = 0.350). These levels were lower than adults in EVC (*M* = 0.208, permutation *p* < 0.001), LOC (*M* = 0.224, *p* < 0.001), precuneus (*M* = 0.087, *p* = 0.004), and EAC (*M* = 0.068, *p* = 0.007); all other regions did not differ between groups (AG: *M* = 0.015, *p* = 0.655; PCC: *M* = 0.056, *p* = 0.137; mPFC: *M* = -0.020, *p* = 0.527; hippocampus: *M* = -0.012, *p* = 0.669). In sum, there is strong evidence of a shared response across infants, not just in visual regions, but also in regions involved in narrative processing in adults.

### Flattened hierarchical gradient of event timescales in the infant brain

Given that infants process the movie in a similar way to each other, we next asked whether their neural activity contains evidence of event segmentation, as in adults. Our analysis tested whether infant brains transition through discrete event states characterized by stable voxel activity patterns, which then shift into new stable activity patterns at event boundaries. We used a computational model to characterize the stable neural event patterns of infant and adult brains ^[25]^. We analyzed the data from infant and adult groups separately. Within each group, we repeatedly split the data in half, with one set of participants forming a training set and the other a test set. We learned the model on the training set using a range of event numbers from 2 to 21, and then applied it to the test set. Model fit was assessed by the log probability of the test data according to the learned event segmentation (i.e., log-likelihood). In a searchlight analysis, we assigned to each voxel the number of events that maximized the log-likelihood of the model for activity pattern from surrounding voxels across test iterations. In an ROI analysis, the patterns were defined from all voxels in each ROI.

In adults, we replicated previous work showing a hierarchical gradient of event timescales across cortex, with more/shorter events in early visual compared to higher-order associative regions (Figure 2A). Qualitative inspection revealed that boundaries in EVC seemed to correspond to multiple types of visual changes in the movie (e.g., between camera angles, viewpoints of main characters), while precuneus boundaries included major plot points (e.g., arrival of pilot, flying machine breaking, return of blueprint that fixes machine). In contrast, infants did not show strong evidence of a hierarchical gradient. In fact, the model performed optimally with fewer/longer events across the brain (Figure 2B). Although the timing of infant EVC boundaries differed from adults, the infant precuneus boundary was part of the set of adult precuneus boundaries (corresponding to the flying machine breaking). These findings support the third hypothesis of infants being biased to longer event timescales throughout the cortical hierarchy.

**Figure 2.**
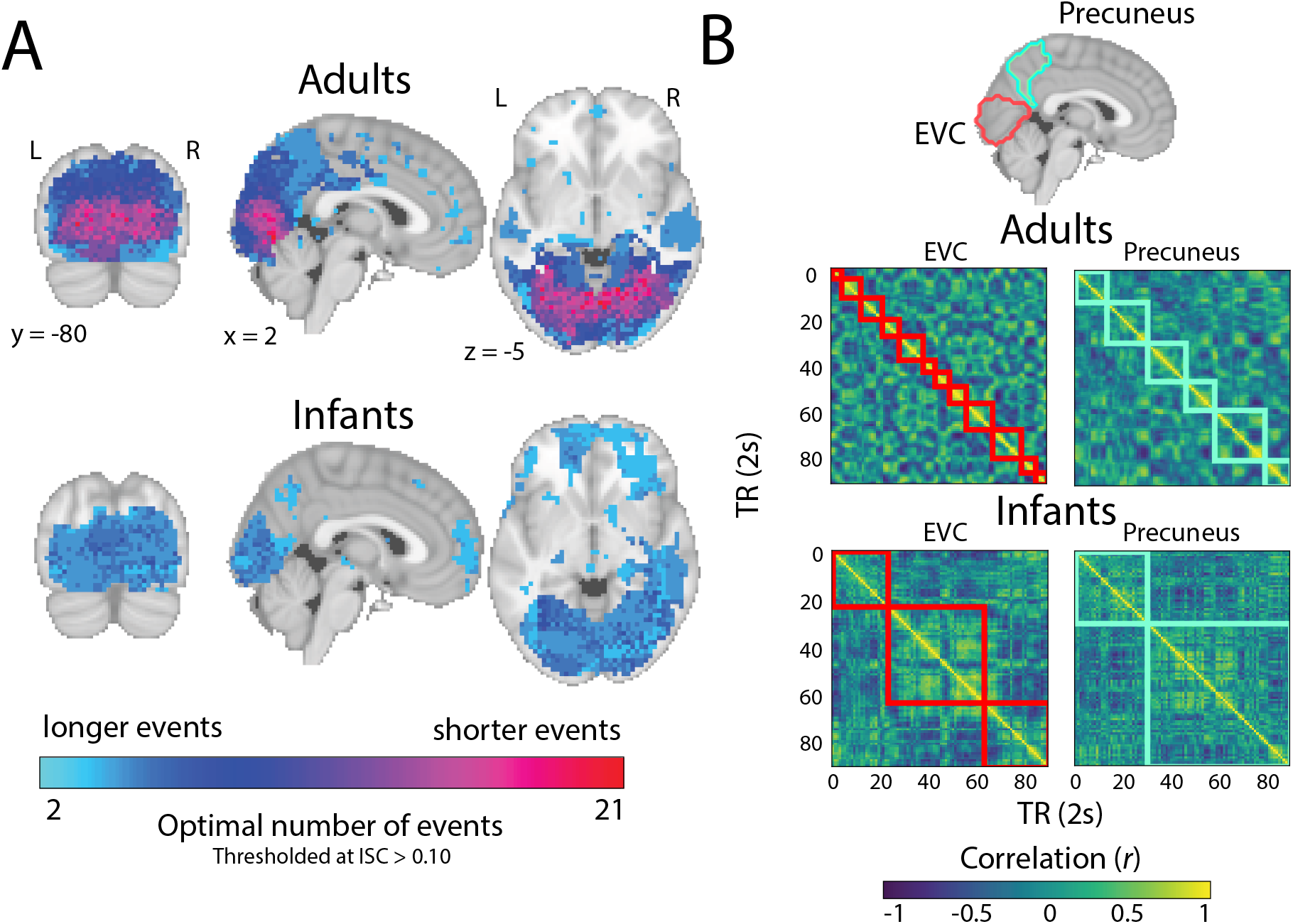
Event structure across the adult and infant brain. (A) The optimal number of events for a given voxel was determined via a searchlight across the brain, which found the number of events that maximized the model log-likelihood in held-out data. Voxels with an average ISC value greater than 0.10 are plotted for visualization purposes. In adults, there was a clear difference in the number of events found in early visual regions vs. higher-order associative regions, but this was not present in infants. Instead, there was a flattened hierarchy in the infant brain, with fewer/longer events in both early visual and associative regions. (B) Example timepoint-by-timepoint correlation matrices in EVC and precuneus in the two groups. The model event boundaries found for each age group are outlined in red (EVC) and aqua (precuneus).

### Coarser but reliable event structure across brain regions in infants

The above analysis provides a qualitative description of the timescale of event processing in the infant brain. However, comparing the *relative* model fits for different timescales does not allow us to assess whether the model fit at the optimal timescale is significantly above chance. To quantify whether these learned events truly demarcated state changes in neural activity patterns, we used nested cross-validation. For each ROI, we followed the steps above for finding the optimal number of events, but critically, held one participant out of the analysis completely (and iterated so each participant was held out once). On each leave-one-participant-out iteration, the number of optimal events in the remaining N-1 training participants could vary; the held-out participant had no impact on the learned event model. The model with the optimal event structure in the training participants was then fit to the held-out participant’s data. To establish a null distribution, we rotated the held-out participant’s data in time. A *z*-score of the log-likelihood for the actual result versus the null distribution was calculated to determine whether the learned event structure generalized to a new participant better than chance (Figure 3A). This analysis can tell us whether the smaller number of events observed in infants reflects true differences in processing granularity between adults and infants, or results from combining across infants who have idiosyncratic event structures.

**Figure 3.**
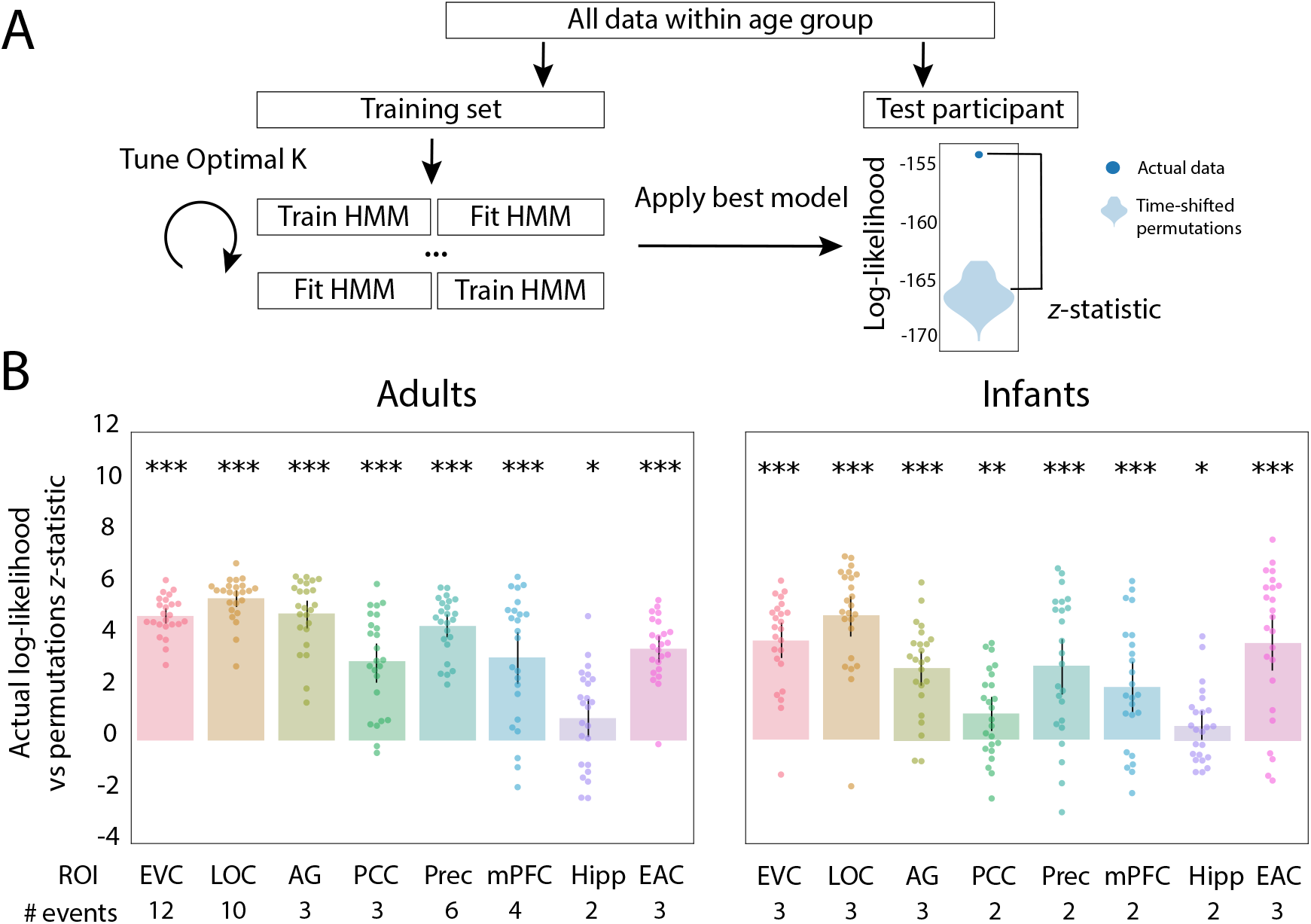
Nested cross-validation of adult and infant age groups. (A) Schematic explaining the nested cross-validation procedure for computing the reliability of event segmentation. (B) Across ROIs and for both adults and infants, the model fit real better than permuted held-out data. The number of events that optimized model log-likelihood in the full sample of participants is labeled below the x-axis. Dots represent individual participants and error bars represent 95% CIs of the mean from bootstrap resampling. *** *p* < 0.001, ** *p* < 0.01, * *p* < 0.05, ∼*p* < 0.1.

Overall, the models for different ROIs reliably fit independent data (Figure 3B). All ROIs were significant in adults (EVC: *M* = 4.85, CI = [4.55, 5.16], *p* < 0.001; LOC: *M* = 5.53, CI = [5.18, 5.86], *p* < 0.001; AG: *M* = 4.94, CI = [4.38, 5.43], *p* < 0.001; PCC: *M* = 3.07, CI = [2.27, 3.77], *p* < 0.001; precuneus: *M* = 4.46, CI = [4.02, 4.91], *p* < 0.001; mPFC: *M* = 3.23, CI = [2.21, 4.20], *p* < 0.001; hippocampus: *M* = 0.907, CI = [0.147, 1.64], *p* = 0.014; EAC: *M* = 3.60, CI = [3.06, 4.06], *p* < 0.001). This was also true for infants (EVC: *M* = 3.89, CI = [3.20, 4.57], *p* < 0.001; LOC: *M* = 4.87, CI = [4.08, 5.62], *p* < 0.001; AG: *M* = 2.83, CI = [2.15, 3.49], *p* < 0.001; PCC: *M* = 1.05, CI = [0.394, 1.70], *p* = 0.002; precuneus: *M* = 2.91, CI = [1.81, 3.95], *p* < 0.001; mPFC: *M* = 2.08, CI = [1.09, 3.07], *p* < 0.001; hippocampus: *M* = 0.567, CI = [0.006, 1.17], *p* = 0.046; EAC: *M* = 3.83, CI = [2.72, 4.89], *p* < 0.001). This analysis confirms that the longer event timescales we observed across the cortical hierarchy in infants are reliable.

Infant data are often noisier than adult data ^[35]^, so we took additional steps to verify that the longer events in infants were not a mere byproduct of this noise. In particular, we simulated fMRI data with different levels of noise and fit the event segmentation model at each level. The model tended to over-estimate the optimal number of events (i.e., produce shorter events) as noise increased (Figure S1). This suggests that the smaller number of longer events in infants does not result from increased noise per se.

### Relationship between adult and infant event structure

The optimal number of events for a given region differs across adults and infants, but this does not necessarily mean that the patterns of neural activity are unrelated. The coarser event structure in infants may still be present in the adult brain, with their additional events carving up these longer events more finely. Conversely, the finer event structure in adults may still be developing in the infant brain, such that it may be present but have a less optimal fit. We thus investigated whether event structure from one group could explain the neural activity of individual participants in the other group (Figure 4). We compared this across-group prediction to a baseline of how well other members of the same group could explain an individual’s neural activity. If event structure better explains neural data from the same age group compared to the other age group, then we can conclude that event structures differ between age groups.

**Figure 4.**
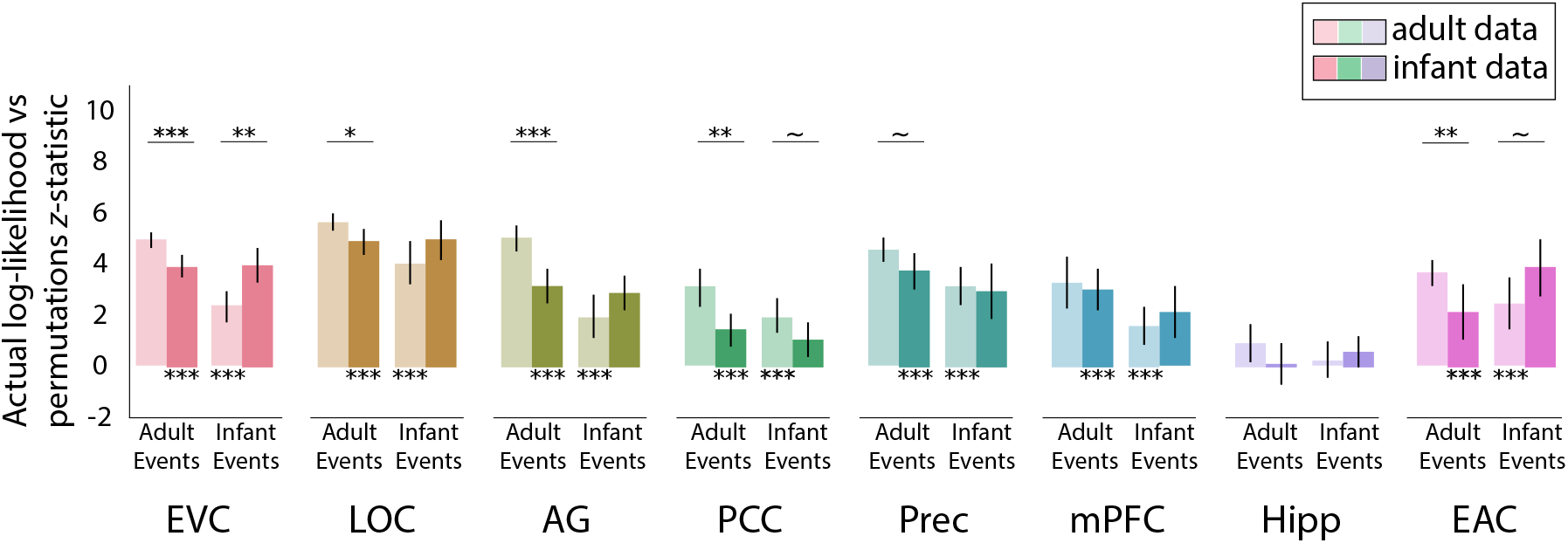
Reliability of event structure for models learned on participants of the same vs. other age group. (A) Light bars indicate fit of adult and infant event structures to adult data, and dark bars indicate fit of adult and infant event structures to infant data. Note that the fits to the same group (adult events in adults, infant events in infant) are simply replotted from Figure 3, without duplicating the statistics. Overall, event structures learned from adults and infants fit data from the other group (clearest in EVC and LOC). However, in several regions, these fits were weaker than to data from the same group (clearest in EVC, LOC, AG, PCC, and EAC for adult events, and in EVC and EAC for infant events). Error bars represent 95% CIs of the mean from bootstrap resampling. *** *p* < 0.001, ** *p* < 0.01, * *p* < 0.05, ∼*p* < 0.1.

When event segmentation models fit to adults were applied to infant neural activity, all ROIs except hippocampus showed significant model fit over permutations (EVC: *M* = 3.81, CI = [3.40, 4.26], *p* < 0.001; LOC: *M* = 4.82, CI = [4.30, 5.29], *p* < 0.001; AG: *M* = 3.12, CI = [2.45, 3.72], *p* < 0.001; PCC: *M* = 1.43, CI = [0.796, 2.03], *p* < 0.001; precuneus: *M* = 3.68, CI = [2.93, 4.34], *p* < 0.001; mPFC: *M* = 2.95, CI = [2.18, 3.73], *p* < 0.001; hippocampus: *M* = 0.106, CI = [-0.659, 0.892], *p* = 0.794; EAC: *M* = 2.12, CI = [1.03, 3.18], *p* < 0.001). This suggests that although infants and adults had a different optimal number of events in these regions, there was some overlap in their event representations. In most of these regions, models trained on adults showed significantly or marginally better fit to adults than to infants, (EVC: *M* = 1.04, CI = [0.490, 1.56], *p* < 0.001; LOC: *M* = 0.715, CI = [0.152, 1.31], *p* = 0.012; AG: *M* = 1.82, CI = [0.952, 2.65], *p* < 0.001; PCC: *M* = 1.64, CI = [0.691, 2.68], *p* = 0.002; precuneus: *M* = 0.781, CI = [-0.024, 1.61], *p* = 0.054; EAC: *M* = 1.48, CI = [0.385, 2.65], *p* = 0.008), suggesting that adult-like event structure is still developing in these regions. Indeed, how well adult event structure fit an infant was related to infant age in LOC (*r* = 0.457, *p* = 0.026). No other ROIs showed a reliable relationship with age (*p*s *>* 0.10), though we had a relatively small sample and truncated age range for evaluating individual differences.

When event segmentation models learned from infants were applied to adult neural activity, all ROIs except hippocampus showed significant model fit over permutations (EVC: *M* = 2.33, CI = [1.73, 2.89], *p* < 0.001; LOC: *M* = 3.99, CI = [3.14, 4.81], *p* < 0.001; AG: *M* = 1.90, CI = [1.08, 2.75], *p* < 0.001; PCC: *M* = 1.93, CI = [1.33, 2.63], *p* < 0.001; precuneus: *M* = 3.08, CI = [2.35, 3.84], *p* < 0.001; mPFC: *M* = 1.59, CI = [0.846, 2.31], *p* = 0.001; hippocampus: *M* = 0.215, CI = [-0.433, 0.941], *p* = 0.584; EAC: *M* = 2.41, CI = [1.46, 3.42], *p* < 0.001). Infant event models explained infant data better than adult data only in EVC (*M* = 1.56, CI = [0.585, 2.42], *p* = 0.006) and marginally in EAC (*M* = 1.42, CI = [-0.106, 2.85], *p* = 0.058). Intriguingly, infant event models showed marginally better fit to *adult* vs. infant neural activity in PCC (*M* = -0.879, CI = [-1.84, 0.126], *p* = 0.078).

Altogether, the finding that events from one age group significantly fit data from the other age group shows that infant and adult event representations are related in all regions except the hippocampus. Nonethe-less, the better fit in some ROIs when applying events from one age group to neural activity from the same vs. other age group provides evidence that these event representations are not the same and may in fact change over development, compatible with the second and third hypotheses that the infant brain segments experience differently than adults.

### Expression of behavioral event boundaries

We took a brain-based, data-driven approach to discovering event representations in adults and infants, but how do these neural event representations relate to behavior? The behavioral task of asking participants to explicitly parse or annotate a movie is not possible in infants. However, in adults such annotations of high-level scene changes have been shown to align with neural event boundaries in the adult AG, precuneus, and PCC^[25]^. Given that we found that adult neural event boundaries in these same regions significantly fit infant data, we tested whether adult behavioral event boundaries might also be reflected in the infant brain.

We collected behavioral event segmentation data from 22 independent adult participants who watched Aeronaut while identifying salient boundaries^[24]^ (Figure 5A). Participants were not instructed to annotate at any particular timescale, and were simply asked to indicate when it felt like a new event occurred. We quantified the fit of behavioral boundaries to neural activity by calculating the difference in pattern similarity between two timepoints within vs. across boundaries, equating temporal distance. Results were weighted by the number of unique timepoint pairs that made up the smaller group of correlations (e.g., close to the boundary, there are fewer across event pairs than within event pairs). A more conservative approach considered timepoint pairs within vs. across event boundaries anchored to the same timepoint (Figure S2). To the extent that behavioral segmentation aligned with event boundaries in a region, we expected greater neural similarity for timepoints within events.

**Figure 5.**
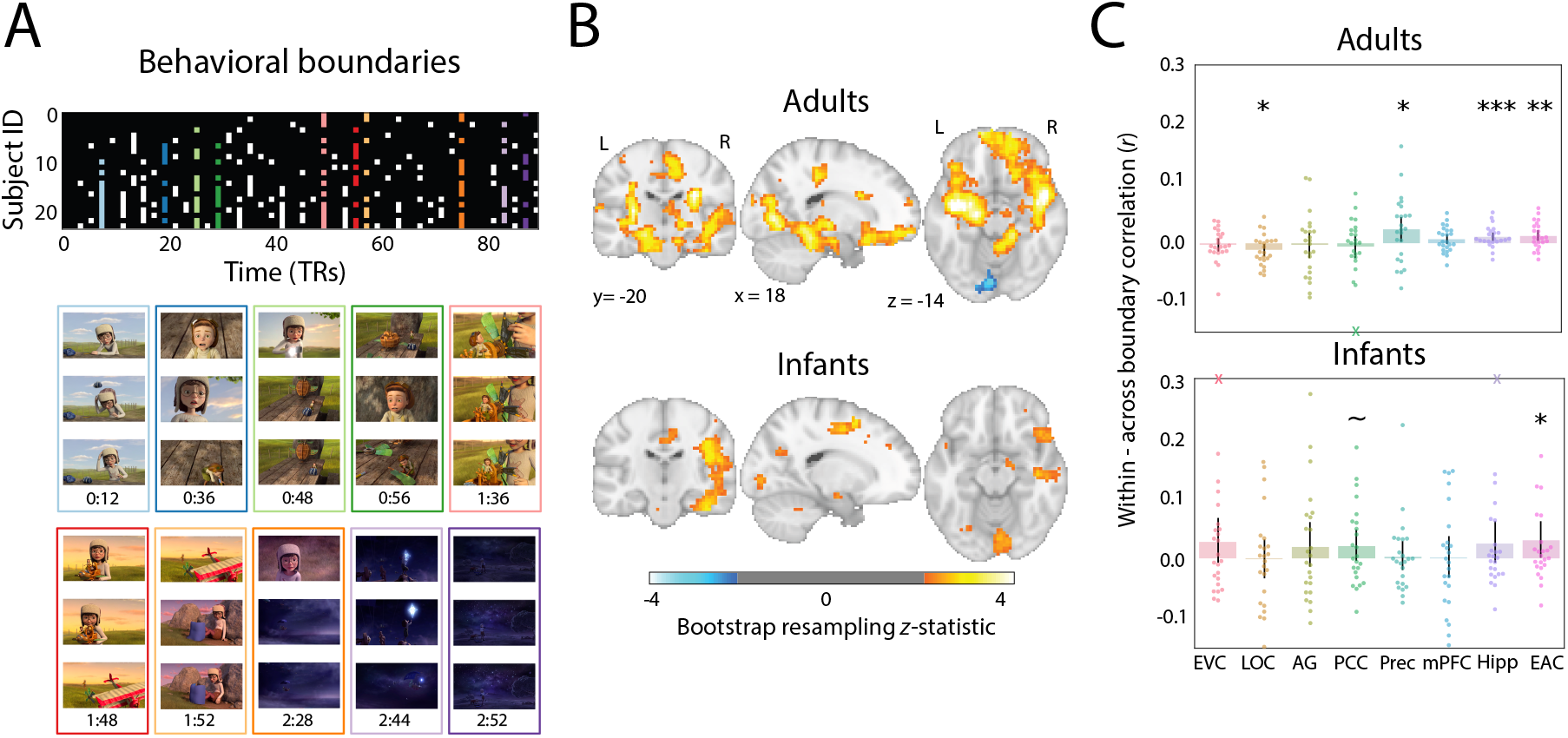
Relating behavioral boundaries to neural activity. (A) Matrix showing which behavioral participants indicated the presence of an event boundary at each TR in the movie. The 10 TRs with the highest percentage of agreement robust to response time adjustment were used as event boundaries (colored columns; see Methods). Movie frames from the TR before, during, and after each event boundaries are depicted below for qualitative inspection. (B) Whole-brain searchlight analysis for each age group comparing pattern similarity between timepoints drawn from within vs. across behavioral event boundaries. Bootstrapped *z*-scores are thresholded at *p* < 0.05, uncorrected. (C) ROI analysis of difference in pattern similarity within minus between behavioral events. Dots represent individual participants and error bars represent 95% CIs of the mean from bootstrap resampling. One adult participant with a value beyond the y-axis range for PCC is indicated with an X at the negative edge. Infant participants with values beyond the y-axis range for EVC and hipppocampus are indicated with Xs at the positive edge. *** *p* < 0.001, ** *p* < 0.01, * *p* < 0.05, ∼*p* < 0.1.

We performed this analysis for both whole-brain searchlights and ROIs. In adults, the searchlight revealed greater pattern similarity within vs. across behavioral event boundaries throughout the brain, including visual regions, medial frontal cortex, bilateral hippocampus, supramarginal gyrus, and posterior cingulate (Figure 5B). This generally fits with previous work showing that neural event in visual and semantic regions can relate to behavioral boundaries^[25]^. For the ROIs (Figure 5C), we found significantly greater pattern similarity within vs. across behavioral boundaries in precuneus (*M* = 0.024, CI = [0.003, 0.045], *p* = 0.024), hippocampus (*M* = 0.011, CI = [0.004, 0.018], *p* < 0.001), and EAC (*M* = 0.013, CI = [0.004, 0.022], *p* = 0.002), as well as significantly *less* pattern similarity within vs. across behavioral boundaries in LOC (*M* = -0.011, CI = [-0.022, -0.001], *p* = 0.028); the other regions were insensitive to the behavioral boundaries (EVC: *M* = -0.002, CI = [-0.013, 0.008], *p* = 0.622; AG: *M* = -0.003, CI = [-0.025, 0.018], *p* = 0.742; PCC: *M* = -0.005, CI = [-0.025, 0.013], *p* = 0.602; mPFC: *M* = 0.007, CI = [-0.002, 0.015], *p* = 0.108). Given that most of the non-significant ROIs in this analysis showed reliable event segmentation overall (Figure 3B), the behavioral event boundaries may have been misaligned, for example by anticipation^[49]^, or focused at a particular timescale that could have been modified through instructions.

In infants, several regions showed greater pattern similarity within vs. across behavioral boundaries in the searchlight analysis, including visual regions, supramarginal gyrus, posterior and anterior cingulate, and right lateral frontal cortex. In the ROIs, there was significant greater pattern similarity within vs. across behavioral boundaries in EAC (*M* = 0.030, CI = [0.001, 0.062], *p* = 0.044) and marginal in PCC (*M* = 0.021, CI = [-0.003, 0.049], *p* = 0.088); the other regions were insensitive to the behavioral boundaries (EVC: *M* = 0.027, CI = [-0.007, 0.068, *p* = 0.104; LOC: *M* = -0.001, CI = [-0.033, 0.031], *p* = 0.894); AG: *M* = 0.019, CI = [-0.014, 0.060], *p* = 0.306); precuneus: *M* = 0.003, CI = [-0.019, 0.029], *p* = 0.878; mPFC: *M* = 0.000, CI = [-0.032, 0.037], *p* = 0.994; hippocampus: *M* = 0.025, CI = [-0.006, 0.062], *p* = 0.144). Alignment between behavioral boundaries and event structure in EAC was also related to infants age (*r* = 0.326, *p* = 0.036). This is especially striking because adult EAC was also aligned with adult behavioral segmentation. Thus, infants can have neural representations related to how adults explicitly segment a movie, long before they can perform the behavior, understand task instructions, or even speak. Nonetheless, this occurred in a small number of regions that did not fully overlap with adults, suggesting functional changes over development in the behavioral relevance of neural signals for event segmentation.

### Event structure in an additional infant cohort

To provide additional evidence of coarser event structure in infancy, we applied our analyses to a more heterogeneous sample of infants who watched a different, short cartoon movie (“Mickey”) during breaks between tasks for other studies. We first asked whether there were consistent neural responses within a group of 15 adults, and found a similar topography of ISC as in the main cohort, with significant values in EVC, LOC, AG, PCC, precuneus, and EAC (Figure S3). ISC was weaker in 15 infants, though still significant in EVC and LOC. Weaker ISC may potentially be related to the broader age range of the infants (4–33 months) – almost two additional years – given the dramatic developmental changes that occur in this age range and the reliance of ISC on common signal across participants. The smaller sample size, smaller stimulus display (1/4 size), and different movie may also have been factors that influenced ISC. We next fit the event segmentation model and found a hierarchical gradient of event timescales across adult cortex, with more events in sensory regions and fewer events in associative regions (Figure S4). However, this gradient was less pronounced than in the Aeronaut dataset, with fewer events in visual regions. In all but mPFC and hippocampus, event structure significantly explained held-out adult data. We therefore found a similar pattern of results in adults across both datasets. In infants, there was no evidence of a hierarchical gradient in the number/length of events (Figure S4). The model again favored fewer/longer events across regions, and these events reliably fit neural activity from a held-out participant in EVC, LOC, precuneus, mPFC, and hippocampus (Figure S4). This additional sample provides further evidence for the third hypothesis that there is reliable but coarser event structure across the infant brain.

## Discussion

In this study, we investigated neural event segmentation using a data-driven, computational approach in adults and infants watching a short movie. We found synchronous processing of the movie and reliable event structure in both groups. In adults, we replicated a previously observed hierarchical gradient of timescales in event processing across brain regions. However, this gradient was flattened in infants, who instead had coarse neural event structure across regions. Although event structure from one age group provided a reliable fit to the other age group, suggesting some similarity in their representations, adult event structure best fit adult data, suggesting developmental differences. Furthermore, behavioral boundaries aligned with event structure in overlapping but distinct regions across the two age groups. Altogether, this study provides novel insights into how infants represent continuous experience, namely that they automatically segment experience into discrete events, as in adults, but at a coarser granularity lacking a hierarchical gradient.

Different mechanisms could be responsible for longer event timescales in infant visual regions. One account could involve which inputs are processed by the infant brain. Developmental differences in sensation (e.g., acuity), perception (e.g., object recognition), and/or attention (e.g., selection, vigilance) could limit the number of visual features or transients that are registered, decreasing the number of events that could be represented. By this account, the architecture for hierarchical event processing may be intact in infancy but not fully activated given input constraints. An alternative account is that the infant brain receives rich input but that the architecture itself takes time to develop. This is consistent with findings that infants integrate sensory inputs more flexibly in time. Namely, young children bind multisensory and visual features over longer temporal receptive windows than older children and adults^[44–47]^. Interestingly, this diminished temporal precision may be advantageous to infants when gathering information about objects, labels, and events in their environment ^[50]^. For instance, infants may better extract meaning from social interactions if they can bind together continuously unfolding visual, auditory, and emotional information; accordingly, toddlers with autism spectrum disorder have shorter than normal temporal receptive windows ^[45]^. This behavioral literature has been agnostic to how or why temporal receptive windows are dilated in infancy, but the lack of neural gradient may be related. Future work combining behavioral and neural approaches to temporal processing and attention could inform this relationship.

A less theoretically interesting explanation for the smaller number of events in infant visual regions could be model bias. For example, the model may default to fewer events in heterogeneous participant groups. Although the Aeronaut dataset had a relatively narrow band of absolute age (9 months), there are dramatic cognitive and neural changes during the first year of life ^[51]^. We found only limited evidence for developmental trajectories in infant event representation (e.g., in how well adult event structure fit infant LOC). That said, to test whether heterogeneity and noise reduce the estimated number of events, we performed a series of simulations. Contrary to what would be predicted, the optimal number of events *increased* when we added more noise. This is inconsistent with attributing the fewer/longer events in infant visual regions to greater functional and anatomical variability. Nonetheless, research with larger samples targeted at more specific age bands across a broader range of development could inform whether there are meaningful changes in event structure during infancy.

In contrast to visual regions, higher-order regions of adult and infant brains represented events over a similar timescale. This maturity might be understood in light of the sensitivity of infants to goal-directed actions and events ^[5]^. Young infants both predict the outcomes of actions^[52]^ and are surprised by inefficient paths towards a goal^[53]^ when a causal agent is involved. Unambiguous agency also increases the ability of older infants to learn statistical structure^[54]^, suggesting that the infant mind may prioritize agency. Indeed, infants are better at imitating a sequence of actions that have hierarchical versus arbitrary structure^[3,55]^ and show better memory for events that have a clear agent^[56]^, perhaps because of a propensity to segment events according to goals during encoding. Our findings of seemingly mature temporal processing windows in higher-order but not visual regions in infancy suggests that coarser event representations may precede fine-grained representations. Longitudinal tracking of infants’ neural event representations could inform this possibility.

We chose the Aeronaut movie for this study because it was dynamic, had appropriate content for infants, and completed a narrative arc within a short timeframe. Nevertheless, the event boundaries detected in the brain and behavior surely depended in part on the particularities of this movie, including both low-level changes such as color and motion and high-level changes to characters and locations. We view this as a first step, parallel to initial adult studies^[25]^, in demonstrating the existence and general characteristics of neural event segmentation in infants. Relying on an off-the-shelf movie makes it more difficult to determine which factors drive segmentation. Adult EVC vs. precuneus boundaries seemed to occur at timepoints that made sense for each region’s respective function (visual changes vs. major plot points), but we recognize that this is a post-hoc, qualitative observation. Nonetheless, our study lays the groundwork for future studies to manipulate video content and test various factors. For instance, in adults it was subsequently determined that neural event patterns in mPFC reflect event schemas^[26]^ and neural event patterns in precuneus and mPFC track surprisal with respect to beliefs^[57]^. Future infant studies could experimentally manipulate the content and timescales of dynamic visual stimuli to better understand the nature of longer neural event structure in infants.

The nature of conducting fMRI research in awake infants means our study has several limitations. First, there was more missing data in the infant group than in the adult group from eye closure and eye movements. We partially addressed this issue by introducing a new variance parameter to the computational model, but acknowledge that it remains an unavoidable difference in the data. Second, our analyses were conducted in a common adult standard space, requiring alignment across participants. Because of uncertainty in the localization and extent of these regions in infants, we defined our ROIs liberally. This may explain the curious finding of reliable event structure and relation to behavioral boundaries in EAC for both adults and infants. Indeed, the EAC ROI encompassed secondary auditory regions and superior temporal cortex, which is important for social cognition, motion, and face processing^[58,59]^, and shows a modality-invariant response to narratives^[60]^. Future work could define ROIs based on a child atlas^[61]^, although that would complicate comparison to adults. Alternatively, ROIs could be defined in each individual using a functional localizer task, though collecting both movie and localizer data from a single awake infant session is difficult. Regardless, in other studies we have successfully used adult-defined ROIs^[35,36]^. Finally, the age range of infants was wide from a developmental perspective. This was a practical reality given the difficulty of recruiting and testing awake infants in fMRI. Nevertheless, it is potentially problematic in that our model can only capture event structure shared across participants. Future work could focus on larger samples from narrower age bands to ascertain how changes in neural event representations relate to other developmental processes (e.g., language acquisition, motor development).

In conclusion, we found that infants automatically segment continuous experience into discrete neural events, but do so in a coarser way than the corresponding brain regions in adults and without a resulting hierarchical gradient in the timescale of event processing across these regions. This neuroscientific approach for accessing infant mental representations complements a rich body of prior behavioral work on event cognition, providing a new lens on how infants bring order to the “blooming, buzzing confusion”^[62]^ of their first year.

## Methods and Materials

### Participants

Data were collected from 24 infants aged 3.60–12.70 months (*M* = 7.43, *SD* = 2.70; 12 female) while they watched a silent cartoon (“Aeronaut”). Infants who moved their head excessively (N = 11) or did not look the screen (N = 4) during more than half of the movie were excluded from this total. We further excluded participants when we had to stop the scan less than halfway through the movie because of fussiness or movement (N = 9) and because of technical error (N = 1). For comparison, we also collected data from 24 adult participants aged 18 – 32 years (*M* = 22.54, *SD* = 3.66; 14 female) who watched the same movie. An extraneous adult participant was collected but subsequently excluded to equate infant and adult participant group sizes. Although some infant and adults participants watched Aeronaut one or more additional times in later fMRI sessions, we only analyzed the first session in which we collected usable data in the current paper. The study was approved by the Human Subjects Committee (HSC) at Yale University. All adults provided informed consent, and parents provided informed consent on behalf of their child.

### Materials

Aeronaut is a 3-minute long segment of a short film entitled “Soar” created by Alyce Tzue (https://vimeo.com/148198462). The film was downloaded from YouTube in Fall 2017 and iMovie was used to trim the length. The audio was not played to participants in the scanner. The movie spanned 45.5 visual degrees in width and 22.5 visual degrees in height. In the video, a girl is looking at airplane blueprints when a miniature boy crashes his flying machine onto her workbench. The pilot appears frightened at first, but the girl helps him fix the machine. After a few failed attempts, a blueprint flies into the girl’s shoes, which they use to finally launch the flying machine into the air to join a flotilla of other ships drifting away. In the night sky, the pilot opens his suitcase, revealing a diamond star, and tosses it into the sky. The pilot then looks down at Earth and signals to the girl, who looks up as the night sky fills with stars.

The code used to show the movies on the experimental display is available at https://github.com/ntblab/experiment_menu/tree/Movies/. The code used to perform the data analyses is available at https://github.com/ntblab/infant_neuropipe/tree/EventSeg/; this code builds on tools from the Brain Imaging Analysis Kit (^[63]^; https://brainiak.org/docs/). Raw and preprocessed functional data and anatomical images will be released publicly.

### Data acquisition

Procedures and parameters for collecting MRI data from awake infants were developed and validated in a previous methods paper^[35]^, with key details repeated below. Data were collected at the Brain Imaging Center in the Faculty of Arts and Sciences at Yale University. We used a Siemens Prisma (3T) MRI and the bottom half of the 20-channel head coil. Functional images were acquired with a whole-brain T2* gradientecho EPI sequence (TR = 2s, TE = 30ms, flip angle = 71, matrix = 64×64, slices = 34, resolution = 3mm iso, interleaved slice acquisition). Anatomical images were acquired with a T1 PETRA sequence for infants (TR1 = 3.32ms, TR2 = 2250ms, TE = 0.07ms, flip angle = 6, matrix = 320×320, slices = 320, resolution = 0.94mm iso, radial slices = 30000) and a T1 MPRAGE sequence for adults (TR = 2300ms, TE = 2.96ms, TI = 900ms, flip angle = 9, iPAT = 2, slices = 176, matrix = 256×256, resolution = 1.0mm iso). The adult MPRAGE sequence included the top half of the 20-channel head coil.

### Procedure

Before their first session, infant participants and their parents met with the researchers for a mock scanning session to familiarize them with the scanning environment. Scans were scheduled for a time when the infant was thought to be most comfortable and calm. Infants and their accompanying parents were extensively screened for metal. Three layers of hearing protection (silicon inner ear putty, over-ear adhesive covers, and ear muffs) were applied to the infant participant. They were then placed on the scanner bed on top of a vacuum pillow that comfortably reduced movement. Stimuli were projected directly on to the surface of the bore. A video camera (MRC high-resolution camera) was placed above the participant to record their face during scanning. Adult participants underwent the same procedure with the following exceptions: they did not attend a mock scanning session, hearing protection was only two layers (earplugs and optoacoustics noise-canceling headphones), and they were not given a vacuum pillow. Finally, infants may have participated in additional tasks during their scanning session, whereas adult sessions contained only the movie task (and an anatomical image).

### Gaze coding

Gaze was coded offline by 2-3 coders for infants (*M* = 2.2, *SD* = 0.6) and by 1 coder for adults. Based on recordings from the in-bore camera, coders determined whether the participant’s eyes were on-screen, offscreen (i.e., blinking or looking away), or undetected (i.e., out of the camera’s field of view). In one infant, gaze data were not collected because of technical issues; in this case, the infant was monitored by visual inspection of a researcher and determined to be attentive enough to warrant inclusion. For all other infants, coders were highly reliable: They reported the same response code on an average of 93.2% (*SD* = 5.2%; range across participants = 77.7–99.6%) of frames. The modal response across coders from a moving window of five frames was used to determine the final response for the frame centered in that window. In the case of ties, the response from the previous frame was used. Frames were pooled within TRs, and the average proportion of TRs included was high for both adults (*M* = 98.8%, *SD* = 3.2%; range across participants = 84.4–100%) and infants (*M* = 88.4%, *SD* = 12.2%; range across participants = 55.6–100%).

### Preprocessing

Data from both age groups were preprocessed using a modified FSL FEAT pipeline designed for infant fMRI^[35]^. If infants participated in other tasks during the same functional run (*N* = 12), the movie data were cleaved to create a pseudo-run. Three burn-in volumes were discarded from the beginning of each run/pseudo-run. Motion correction was applied using the centroid volume as the reference — determined by calculating the Euclidean distance between all volumes and choosing the volume that minimized the distance to all other volumes. Slices in each volume were realigned using slice-timing correction. Timepoints with greater than 3mm of translational motion were excluded and temporally interpolated so as not to bias linear detrending. The vast majority of infant timepoints were included after motion exclusion (*M* = 92.6%, *SD* = 9.9%; range across participants = 65.6–100%) and all adult timepoints were included. Timepoints with excessive motion and timepoints during which eyes were closed for a majority of frames in the volume were excluded from subsequent analyses. The signal-to-fluctuating-noise ratio (SFNR) was calculated^[64]^ and thresholded to form the mask of brain vs. non-brain voxels. Data were spatially smoothed with a Gaussian kernel (5mm FWHM) and linearly detrended in time. AFNI’s (https://afni.nimh.nih.gov) despiking algorithm was used to attenuate aberrant timepoints within voxels. After removing excess burn-out TRs, functional data were *z*-scored within run.

The centroid functional volume was registered to the anatomical image. Initial alignment was performed using FLIRT with 6 degrees of freedom (DOF) and a normalized mutual information cost function. This automatic registration was manually inspected and then corrected if necessary using mrAlign from mrTools (Gardener lab). To compare across participants, functional data were further transformed into standard space. For infants, anatomical images were first aligned automatically (FLIRT) and then manually (Freeview) to an age-specific MNI infant template ^[65]^. This infant template was then aligned to adult MNI standard (MNI152). Adult anatomical images were directly aligned to the adult MNI standard. For all analyses, we only considered voxels included in the intersection of all infant and adult brain masks.

In an additional exploratory analysis, we re-aligned participants’ anatomical data to the adult standard using ANTs ^[66]^, a non-linear alignment algorithm. For infants, an initial linear alignment with 12 DOF was used to align anatomical data to the age-specific infant template, followed by non-linear warping using diffeomorphic symmetric normalization. Then, as before, we used a predefined transformation (12 DOF) to linearly align between the infant template and adult standard. For adults, we used the same alignment procedure, except participants were directly aligned to adult standard. Results using this non-linear procedure were nearly identical to the original analyses (Figure S5).

### Regions of interest

We performed analyses over the whole brain and in regions of interest (ROIs). We defined the ROIs using the Harvard-Oxford probabilistic atlas^[67]^ (0% probability threshold) in early visual cortex (EVC), lateral occipital cortex (LOC), angular gyrus (AG), precuneus, early auditory cortex (EAC), and the hippocampus. We used functionally defined parcellations obtained in resting state^[68]^ to define two additional ROIs: medial prefrontal cortex (mPFC) and posterior cingulate cortex (PCC). We included these regions because of their involvement in longer time-scale narratives, events, and integration ^[69]^.

### Intersubject correlation

We assessed whether participants were processing the movie in a similar way using intersubject correlation (ISC^[39,70]^). For each voxel, we correlated the timecourse of activity between a single held-out participant and the average timecourse of all other participants in a given age group. We iterated through each participant and then created the average ISC map by first Fisher-transforming the Pearson correlations, averaging these transformed values, and then performing an inverse Fisher-transformation on the average. We visualize the whole-brain map of the intersubject correlations for adults and infants separately, thresholded at a correlation of 0.10.

For the ROI analysis, the voxel ISCs within a region were averaged separately for each held-out participant using the Fisher-transform method described above. Statistical significance was determined by bootstrap resampling. We randomly sampled participants with replacement 1,000 times, on each iteration forming a new sample of the same size as the original group, then averaged their ISC values to form a sampling distribution. The *p*-value was calculated as the proportion of resampling iterations on which the group average had the opposite sign as the original effect, doubled to make it two-tailed. For comparing ISC across infant and adult groups, we permuted the age group labels 1,000 times, each time recalculating ISC values for these shuffled groups and then finding the difference of group means. This created a null distribution for the difference between age groups.

### Event segmentation model

To determine the characteristic patterns of event states and their structure, we applied a Hidden Markov Model (HMM) variant^[25]^ available in BrainIAK ^[63]^ to the average fMRI activity of participants from the same age group. This model uses an algorithm that alternates between estimating two related components of stable neural events: (1) multivariate event patterns and (2) their event structure (i.e., placement of boundaries between events). The constraints of the model are that each event state is only visited once, and that staying versus transitioning into a new event state have the same prior probability. Model fitting stopped when the log probability that the data were generated from the learned event structure (i.e., log-likelihood ^[71]^) began to decrease.

To deal with missing data in the input (a reality of infant fMRI data), we modified the BrainIAK implementation of the HMM. First, in calculating the probability that each observed timepoint was generated from each possible event model, timepoint variance was scaled by the proportion of participants with data at that timepoint. In other words, if some infants had missing data at a timepoint because of head motion or gaze, the variance at that timepoint was adjusted by the square-root of the maximum number of participants divided by the square-root of the number of participants with data at that point. This meant that even though the model was fit on averaged data that obscured missing timepoints, it had an estimate of the “trustworthiness” of each timepoint. Second, for the case in which missing timepoints persisted after averaging across participants, the log-probability for the missing timepoint was linearly interpolated based on nearby values.

The HMM requires a hyperparameter indicating the number of event states. By testing a range of event numbers and assessing model fit, we determined the optimal number of events for a given voxel or region. We used a cubical searchlight (7×7×7 voxels) to look at the timescales of event segmentation across the whole brain. In a given searchlight, the HMM was fit to the average timecourse of activity for a random split half of participants using a range of event counts between 2 and 21. We capped the maximum number of possible events at 21 to ensure that at least some events would be 3 TRs long. The learned event patterns and structure for each event count were then applied to the average time course of activity for held-out data, and model fit was assessed using the log-likelihood. We iterated through this procedure, each time splitting the data in half differently. The center voxel of the searchlight was assigned the number of events that maximized the average log-likelihood across 24 iterations (chosen to be the same number of iterations as a leave-one-participant-out analysis). This analysis was performed in each searchlight, separately for adults and infants, to obtain a topography of event timescales. We also used this method to determine the optimal number of events for each of our ROIs. In these analyses, the timecourse of activity for every voxel in the ROI was used to learn the event structure.

To test whether a given ROI had statistically significant event structure, we used a nested cross-validation approach. The inner loop of this analysis was identical to what is described above, except that a single participant was completely held out from the analysis. After finding the optimal number of events for all but that held-out participant, the event patterns and structure were fit to that participant’s data. The loglikelihood for those data was compared to a permuted rotation distribution, where the participant’s data were time-shifted for every possible shift value between one and the length of the movie (wrapping around to the beginning). We calculated a *z*-statistic as the difference between the actual log-likelihood and the average log-likelihood of the permuted distribution, divided by the standard deviation of the permuted distribution. We then iterated through all participants and used bootstrap resampling of the *z*-statistics to determine significance. We randomly sampled participants with replacement 1,000 times, on each iteration forming a new sample of the same size as the original group, then averaged their *z*-statistics to form a sampling distribution. The *p*-value was calculated as the proportion of resampling iterations with values less than zero, doubled to make it two-tailed.

### Behavioral segmentation

Behavioral segmentation was collected from 22 naive undergraduate students attending Yale University aged 18 – 22 years (*M* = 18.9, *SD* = 1.0; 14 female). All participants provided informed consent and received course credit. Participants were instructed to watch the Aeronaut movie and press a key on the keyboard when a new, meaningful event occurred. Participants watched a version of the movie with its accompanying audio – a musical track without language. Although the visual input remained the same as the fMRI data collection, these auditory cues may have influenced event segmentation^[72]^. During data collection, participants also evaluated nine other movies, not described here, and verbally recalled each movie after segmenting. We elected to have participants use their own judgement for what constituted an event change. Participants had a 1-minute practice movie to orient them to the task, and the Aeronaut movie appeared in a random order among the list of other movies. To capture “true” event boundaries and avoid contamination by accidental or delayed key presses, we followed a previously published procedure^[24]^. That is, we set a threshold for the number of participants who indicated the same event boundary, such that the number of event boundaries agreed upon by at least that many participants was equal or close to the average number of key presses across participants. Participant responses were binned into 2 TR (4 second) windows. We found 10 event boundaries (11 events) that were each agreed upon by at least 36% of participants and were robust to whether or not key presses were shifted 0.9 seconds to account for response time (for comparison, ∼31% was used in ^[24]^). These event boundaries were then shifted 4 seconds in time to account for the hemodynamic lag.

To evaluate whether these behavioral boundaries predicted neural data, we tested whether voxel activity patterns for timepoints within a boundary were more correlated than timepoints spanning a boundary. This within vs. across boundary comparison has been used previously as a metric of event structure^[25]^. For our analysis, we considered all possible pairs of timepoints within and across boundaries. For each temporal distance from the boundary, we subtracted the average correlation value for pairs of timepoints that were across events from the average correlation value for pairs of timepoints within the same event. At different temporal distances, there are either more or less within-event pairs compared to across-event pairs. To equate the number of within and across event pairs, we subsampled values and recomputed the within vs. across difference score 1,000 times. To combine across distances that had different numbers of possible pairs, we weighted the average difference score for each distance by the number of unique timepoint pairs that made up the smaller group of timepoint pairs (i.e., across-event pairs when temporal distance was low, within-event pairs when temporal distance was high). This was repeated for all participants, resulting in a single weighted within vs. across difference score for each participant. For the ROIs, we used bootstrap resampling of these participant difference scores to determine statistical significance. The *p*-value was the proportion of difference values that were less than zero after 1,000 resamples, doubled to make it two-tailed. For the whole-brain searchlight results, we also used 1,000 bootstrap resamples to determine statistical significance for within vs. across difference scores for each voxel. We then calculated a *z*-score for each voxel as the the distance between the bootstrap distribution and zero, and thresholded the bootstrapped *z*-score map at *p* < 0.05, uncorrected.

## Acknowledgments

We are thankful to the families of infants who participated. We also acknowledge the hard work of the Yale Baby School team, including L. Rait, J. Daniels, and K. Armstrong for recruitment, scheduling, and administration. Thank you to J. Wu, J. Fel, and A. Klein for help with gaze coding and to R. Watts for technical support. We are grateful for internal funding from the Department of Psychology and Faculty of Arts and Sciences at Yale University. N.B.T-B. was further supported by the Canadian Institute for Advanced Research and the James S. McDonnell Foundation (https://doi.org/10.37717/2020-1208).

## Supporting Information

**Figure S1.**
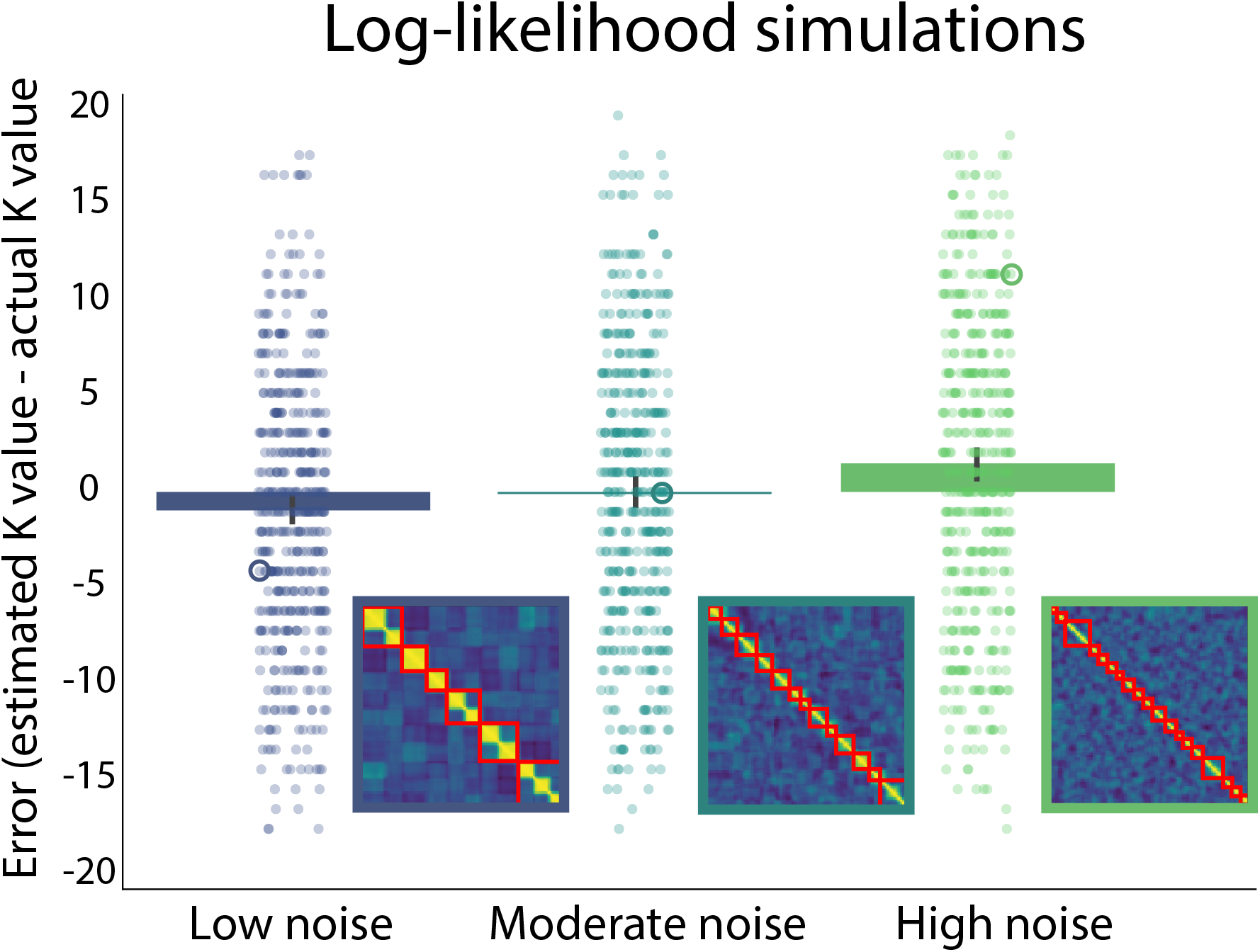
Simulations of log-likelihood metric under different noise regimes. Dots represent differences between the K values that maximized the model log-likelihood and the actual simulated K values for different iterations. One example error value for each noise regime is circled, and an example corresponding timepoint-by-timepoint correlation matrix is inset. Boundaries demarcating the model’s best estimated K value are shown in red. In general, K values were underestimated when noise was low, guessed correctly when noise was moderate, and overestimated when noise was high.

**Figure S2.**
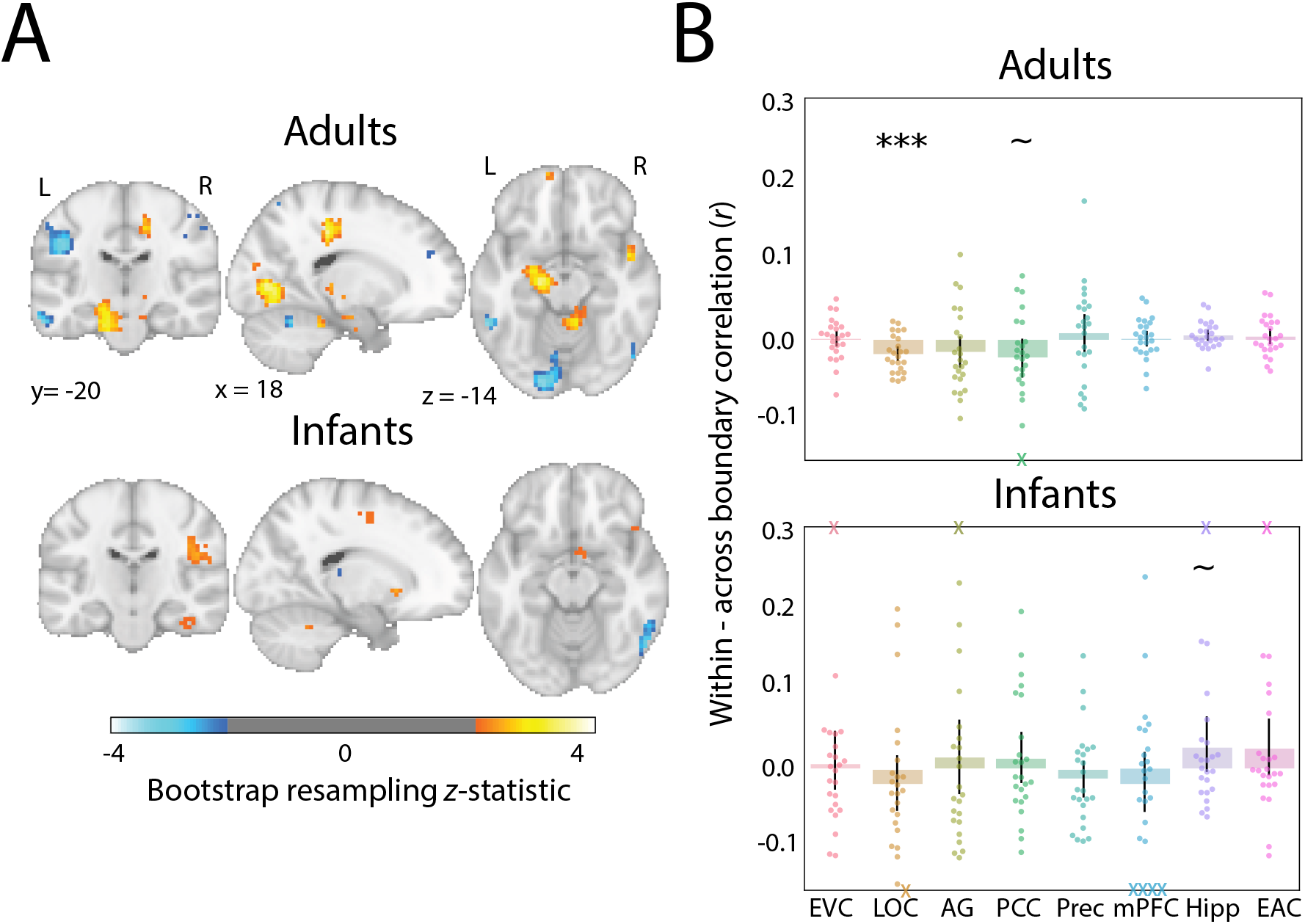
Alternative approach relating behavioral boundaries to neural activity. (A) Whole-brain searchlight analysis for each age group comparing pattern similarity between timepoints drawn from within vs. across behavioral event boundaries when pairs of correlations are anchored to the same timepoint. Bootstrapped *z*-scores are thresholded at *p* < 0.05, uncorrected. (B) ROI analysis of difference in pattern similarity within minus between behavioral events. Dots represent individual participants and error bars represent 95% CIs of the mean from bootstrap resampling. One adult participant with a value beyond the y-axis range for PCC is indicated with an X at the negative edge. Infant participants with values beyond the y-axis range for EVC, AG, hippocampus, and EAC are indicated with Xs at the positive edge, and for LOC and mPFC with Xs at the negative edge. *** *p* < 0.001, ** *p* < 0.01, * *p* < 0.05, ∼*p* < 0.1.

**Figure S3.**
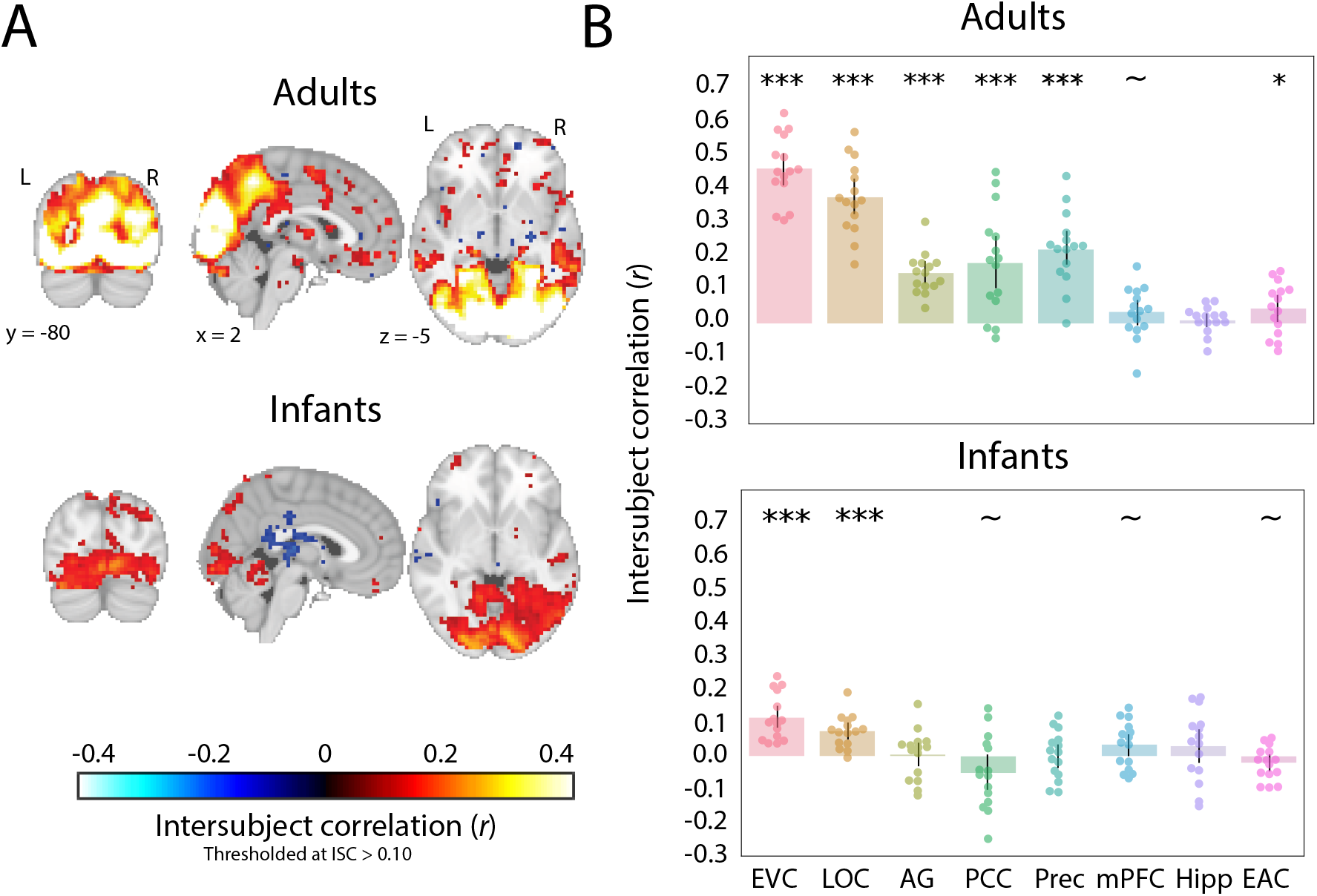
Average leave-one-out intersubject correlation (ISC) in adults and infants watching the Mickey movie. (A) Whole-brain voxel-wise ISC values in the two groups, thresholded for visualization purposes at voxels with correlation values greater than 0.10. (B) ISC values in the ROIs, with the mean at the column height. Dots represent individual participants and error bars represent 95% CIs of the mean from bootstrap resampling. *** *p* < 0.001, ** *p* < 0.01, ∼*p* < 0.1.

**Figure S4.**
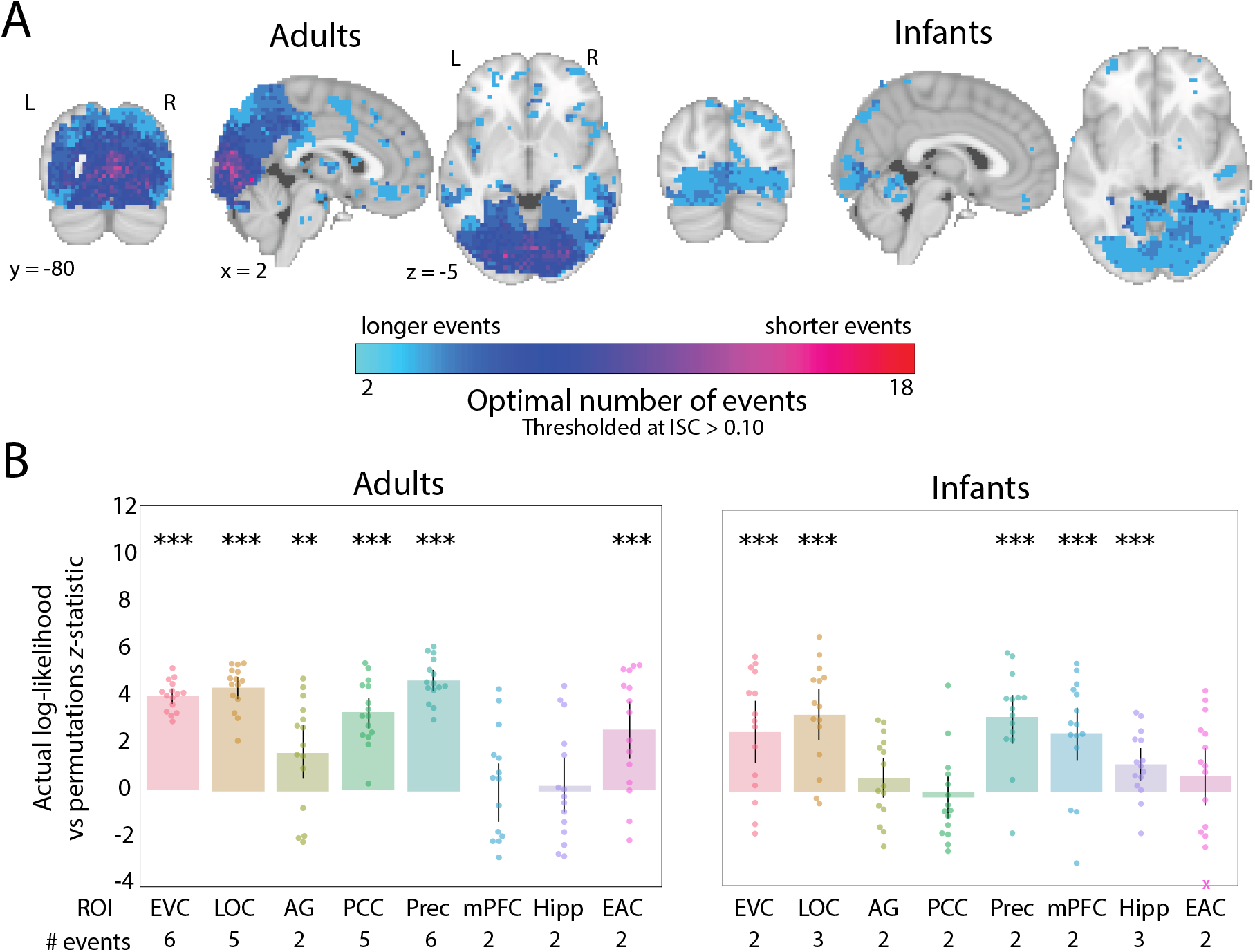
Event structure and reliability for the Mickey movie. (A) The optimal number of events plotted across the brains of adults and infants. Voxels with an average ISC value greater than 0.10 are plotted for visualization purposes. (B) Results of the nested cross-validation procedure for computing the reliability of event segmentation in ROIs for adults and infants, calculated as the *z*-statistic comparing actual and permuted participant data. The number of events that optimized model log-likelihood in the full sample of participants is labeled below the x-axis. Dots represent individual participants and error bars represent 95% CIs of the mean from bootstrap resampling. *** *p* < 0.001, ** *p* < 0.01, * *p* < 0.05.

**Figure S5.**
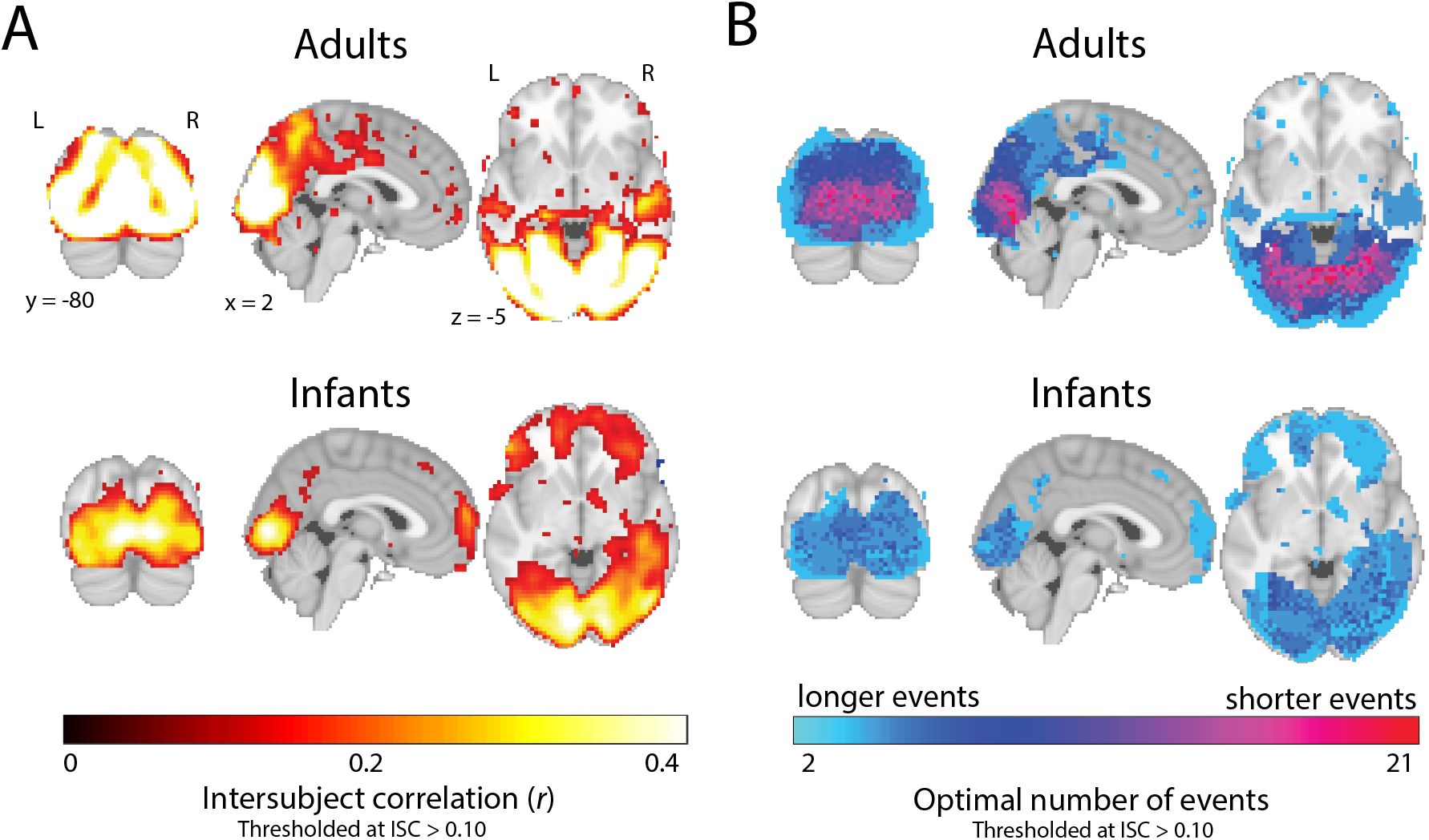
Results from nonlinear anatomical alignment. (A) Whole-brain voxel-wise ISC values in the two groups. (B) The optimal number of events for a given voxel was determined via a searchlight analysis across the brain, which found the number of events that maximized the model log-likelihood in held-out data. In both panels, voxels with an average ISC value greater than 0.10 are plotted for visualization purposes.

## SI Text

### Log-likelihood simulations

To assess whether the log-likelihood metric would be biased to higher or lower numbers of events, we tested how well we could recover event structure in simulated data. We first generated event patterns (voxels by number of events) with values drawn from a standard normal distribution. Because each voxel was treated as an independent source, we used fewer voxels (5) than our actual analyses to better simulate the correlated patterns present in real fMRI data. Event labels were assigned to each of 90 timepoints. We generated 24 “participants” by applying the simulated event patterns to each timepoint with an additional noise component (cov: the covariance matrix of a multivariate normal distribution). The resulting voxel by timepoint matrices were convolved with a double-gamma hemodynamic response function (HRF) using an fMRI simulation package^[73]^ available in BrainIAK ^[63]^. We followed the same analysis approach described in the Methods and Materials section to estimate the optimal number of events. We calculated model error as the difference between the actual simulated number and the estimated optimal number. This was repeated across a range of different possible event numbers (from 2 to 22).

With low noise (cov = 0.1), the timepoint-by-timepoint similarity matrices showed clear block structure along the diagonal. Average error between model estimates and the correct number of events was negative (*M* = -0.931, *p* = 0.003), meaning that the model under-estimated the number of events (Figure S1). When noise increased to a moderate level (cov = 2), model error did not significantly differ from zero (*M* = -0.008, *p* = 0.980), that is, it did not under- or over-estimate the number of events. With high noise (cov = 20), model error was positive (*M* = 1.40, *p* < 0.001), indicating that the model over-estimated the number of events.

### Alternative behavioral boundary approach

We tested whether voxel activity patterns for timepoints within a behavioral boundary were more correlated than timepoints spanning a boundary by considering all possible pairs of timepoints within and across boundaries up to the temporal distance of the largest event. This approach has the advantage of using as many timepoint pairs as possible for calculating within vs. across boundary correlations, but may be vulnerable to increased noise due to comparing timepoints from different parts of the movie. Here we report a more conservative approach for testing how behavioral boundaries predict neural data by only considering timepoint pairs that are equated in temporal distance (as before) but additionally “anchored” to the same timepoint. Namely, for each TR we measured the correlation between the spatial activity pattern at that timepoint and timepoints forward and backward in time at a matched temporal distance. If one timepoint pair was within an event and the other was across an event, we calculated the within minus across boundary correlation. However, if both timepoint pairs were either within an event or across an event, or if one of the timepoint pairs was already included in a different calculation, the within vs. across boundary correlation was not included. We performed this for each temporal distance up to the length of the largest event and calculated the average within vs. across boundary correlation for each subject. For statistical analysis, we used the same bootstrap resampling techniques described in the Methods.

In adults, the searchlight analysis again showed that visual regions and posterior cingulate exhibited significantly greater pattern similarity within vs. across behavioral event boundaries (Figure S2A). In fact, the voxelwise map of the average within vs. across boundary correlation was highly correlated with the map using our main approach (*r* = 0.818). Nonetheless, fewer regions emerged from this more conservative behavioral boundary approach. In infants, we also found several regions showing greater pattern similarity within vs. across behavioral boundaries in infants, where again the voxelwise map was highly correlated with our main results (*r* = 0.893).

In adults, we surprisingly found greater pattern similarity for *across* vs. within timepoint pairs in LOC (*M* = -0.018, CI = [-0.026, -0.008], *p* < 0.001) and marginally in PCC (*M* = -0.022, CI = [-0.048, 0.001], *p* = 0.060); the other regions were insensitive to the behavioral boundaries (EVC: *M* = 0.001, CI = [-0.009, 0.011], *p* = 0.836; AG: *M* = -0.015, CI = [-0.034, 0.004], *p* = 0.114; precuneus: *M* = 0.008, CI = [-0.016, 0.031], *p* = 0.510; mPFC: *M* = 0.002, CI = [-0.008, 0.011], *p* = 0.686); hippocampus: *M* = 0.005, CI = [-0.001, 0.012], *p* = 0.114; EAC: *M* = 0.004, CI = [-0.005, 0.014], *p* = 0.352). In infants, we found marginally greater pattern similarity for within vs. across in the hippocampus (*M* = 0.027, CI = [-0.004, 0.065], *p* = 0.094); the other regions were insensitive to the behavioral boundaries (EVC: *M* = 0.006, CI = [-0.026, 0.048, *p* = 0.750; LOC: *M* = -0.019, CI = [-0.053, 0.016], *p* = 0.342); AG: *M* = 0.014, CI = [-0.031, 0.061], *p* = 0.578); PCC: *M* = 0.012, CI = [-0.018, 0.045], *p* = 0.428; precuneus: *M* = -0.012, CI = [-0.035, 0.010], *p* = 0.320; mPFC: *M* = -0.019, CI = [-0.053, 0.021], *p* = 0.358; EAC: *M* = 0.025, CI = [-0.008, 0.062], *p* = 0.162). Overall, the pattern of results was consistent across the two approaches.

### Mickey dataset

We applied the intersubject correlation and event segmentation analyses to a second, previously collected convenience sample of infants watching a movie between other tasks. Although this dataset has some flaws — fewer participants, much wider age range up to three years, potential for pre-experimental familiarity with the characters, smaller stimulus display, and a mix of scanners — we felt these analyses might provide an initial assessment of generalizability to another movie. fMRI data were collected from 15 sessions with infants aged 4.00 – 32.60 months (*M* = 13.92, *SD* = 8.87; 9 female) watching a silent cartoon lasting 142 s (“Mickey”). This movie was shown on a smaller display than the Aeronaut movie (∼25% of area), spanning 22.75 visual degrees in width and 12.75 visual degrees in height. In this video, a surprise party is thrown where Disney characters dance and play the piano while one character makes an exploding cake in the kitchen. Two infants participated twice after a delay (6.3 months and 2.3 months difference) and were treated as independent sessions. As before, additional infants with head motion above 3 mm (N = 5) or eyes off-screen (N = 2) for more than half of the movie were excluded. We also collected data from 15 adults aged 19 – 27 years (*M* = 21.47, *SD* = 2.90; *N* = 10 female). All adults and 9 infants watched the movie twice in a row within session. For these participants, data were averaged across the two viewings. Infants were collected at the Scully Center at Princeton University (N = 7) and the Magnetic Resonance Research Center (MRRC) at Yale University (N = 8). Adult participants were collected at the Brain Imaging Center (BIC) at Yale University. This study was approved by the Institutional Review Board at Princeton University and the Human Investigation Committee (MRRC) and Human Subjects Committee (BIC) at Yale University. Adults provided informed consented for themselves or their child.

Data acquisition, preprocessing, and analyses were identical to the Aeronaut dataset with two variations: First, infant data were acquired at Princeton using a Siemens Skyra (3T) MRI. Second, functional images for infants were collected under a slightly different functional EPI sequence (TE = 28ms, slices = 36). Adults were collected with the same functional sequence as Aeronaut. Gaze coding was highly reliable: coders reported the same response on an average of 91.3% of frames (*SD* = 5.08%; range = 78.6–98.4%). The average proportion of TRs retained after exclusion for looking off screen was high in adults (*M* = 99.5%, *SD* = 1.01%; range = 96.5–100%) and infants (*M* = 89.0%, *SD* = 13.1%; range = 58.5–100%). Eye-tracking data were not collected for one infant because of experimenter error. Timepoints with less than 3mm of translational motion were included (infants: *M* = 91.3%, *SD* =10.6%, range = 64.8%–100%; adults: 100%).

In the whole-brain analysis, ISC was strongest in visual regions for adults and infants (Figure S3). In the ROI analysis, adult ISC was significant in EVC (*M* = 0.472, CI = [0.423, 0.518], *p* < 0.001), LOC (*M* = 0.384, CI = [0.332, 0.438], *p* < 0.001), AG (*M* = 0.150, CI = [0.121, 0.186], *p* < 0.001), PCC (*M* = 0.188, CI = [0.113, 0.268], *p* < 0.001), precuneus (*M* = 0.228, CI = [0.178, 0.283], *p* < 0.001) and EAC (*M* = 0.045, CI = [0.007, 0.085], *p* = 0.022); marginal in mPFC (*M* = 0.033, CI = [-0.003, 0.068], *p* = 0.070); and not significant in hippocampus (*M* = 0.012, CI = [-0.009, 0.030], *p* = 0.255). Infant ISC was significant in EVC (*M* = 0.117, CI = [0.085, 0.150], *p* < 0.001), LOC (*M* = 0.074, CI = [0.050, 0.099], *p* = 0.001); marginally significant in mPFC (*M* = 0.033 CI = [-0.002, 0.067], *p* = 0.062); marginally significant in the negative direction (likely noise) in PCC (*M* = -0.049, CI = [-0.101, 0.005], *p* = 0.068) and EAC (*M* = -0.022, CI = [-0.048, 0.002], *p* = 0.098); and not significant in AG (*M* = 0.004, CI = [-0.032, 0.038], *p* = 0.865), precuneus (*M* = 0.002, CI = [-0.033, 0.037], *p* = 0.897), or hippocampus (*M* = 0.032, CI = [-0.023, 0.081], *p* = 0.232).

In the searchlight analysis, we applied an HMM to one half of adult or infant participants using a range of event numbers from 2 to 18 and then tested on the second half. This maximum number of events ensured that at least some events were several TRs long, but was less than Aeronaut because Mickey was shorter. Log-likelihood was again used to assess model fit. Similar to our main analyses, visual regions of the adult brain had more events than higher-order regions, although there were fewer events overall (Figure S4). We replicated the flattened gradient of event processing in infants, with the optimal number of events generally low across the brain.

In the nested analysis for reliability of event structure, most ROIs were significant in adults, including EVC (*M* = 3.99, CI = [3.68, 4.32], *p* < 0.001), LOC (*M* = 4.36, CI = [3.86, 4.79], *p* < 0.001), AG (*M* = 1.60, CI = [0.509, 2.80], *p* = 0.008), PCC (*M* = 3.30, CI = [2.61, 3.90], *p* < 0.001), precuneus (*M* = 4.65, CI = [4.19, 5.07], *p* < 0.001), and EAC (*M* = 2.57, CI = [1.37, 3.76], *p* < 0.001); but not mPFC (*M* = 0.007, CI = [-1.31, 1.17], *p* = 0.996) or hippocampus (*M* = 0.222, CI = [-0.902, 1.42], *p* = 0.780). In infants, reliable event structure was found in EVC (*M* = 2.56, CI = [1.23, 3.85], *p* < 0.001), LOC (*M* = 3.31, CI = [2.21, 4.37], *p* < 0.001), precuneus (*M* = 3.21, CI = [2.07, 4.12], *p* < 0.001), mPFC (*M* = 2.49, CI = [1.30, 3.58], *p* < 0.001), and hippocampus (*M* = 1.17, CI = [0.513, 1.86], *p* < 0.001) and not in AG (*M* = 0.585, CI = [-0.228, 1.45], *p* = 0.180), PCC (*M* = -0.267, CI = [-1.15, 0.708], *p* = 0.532) or EAC (*M* = 0.669, CI = [-0.568, 1.84], *p* = 0.260). By generalizing to a different movie, these results provide additional evidence for coarser event representations in infancy.

### Nonlinear alignment

In our main analyses, we used a linear alignment procedure for infant anatomical images (with manual adjustments). However, dramatic developmental differences within and across ages raise the possibility that a nonlinear approach may be more appropriate. We thus used ANTS ^[66]^ to re-align infant and adult brain data from the Aeronaut dataset to adult anatomical data. We repeated the whole-brain ISC analyses and the searchlight analyses of optimal event number. The results were unchanged: ISC was highest in visual regions for both adults and infants (Figure S5A) and there was a gradient in the optimal number of events in adults but not infants (Figure S5B). Thus, our results are robust to these procedures for aligning between infant and adult brains.

